# RNA virus interference via CRISPR/Cas13a system in plants

**DOI:** 10.1101/213645

**Authors:** Rashid Aman, Zahir Ali, Haroon Butt, Ahmed Mahas, Fatima Aljedaani, Muhammad Zuhaib Khan, Shouwei Ding, Magdy Mahfouz

## Abstract

CRISPR/Cas systems confer immunity against invading nucleic acids and phages in bacteria and archaea. CRISPR/Cas13a (known previously as C2c2) is a class 2 type VI-A ribonuclease capable of targeting and cleaving single stranded RNA (ssRNA) molecules of the phage genome. Here, we employ CRISPR/Cas13a to engineer interference with an RNA virus, *Turnip Mosaic Virus* (TuMV), in plants. CRISPR/Cas13a produced interference against green fluorescent protein (GFP) expressing TuMV in transient assays and stable overexpression lines of *Nicotiana benthamiana*. crRNAs targeting the HC-Pro and GFP sequences exhibited better interference than those targeting other regions such as coat protein (CP) sequence. Cas13a can also process pre-crRNAs into functional crRNAs. Our data indicate that CRISPR/Cas13a can be used for engineering interference against RNA viruses, providing a potential novel mechanism for RNA-guided immunity against RNA viruses, and for other RNA manipulations in plants.

## Introduction

CRISPR/Cas adaptive immunity systems confer resistance to invading phages and conjugative plasmids in bacteria and archaea [1–3]. CRISPR/Cas-mediated immunity mechanism involves three steps: adaptation (during which spacers are acquired from the invader genome), processing and biogenesis of crRNAs, and molecular interference against the invader’s genome [4, 5]. CRISPR/Cas systems have been divided into two main classes: class I, in which multi-effector complexes mediate the interference, and class II, which employs single, multi-domain effectors to mediate the interference. These two classes are further subdivided into 6 types and 33 subtypes according to the genomic architecture of the CRISPR array and the signature interference effector. For example, class I includes type I, III, and IV and class II includes type II, V, and VI [6, 7]. Notably, CRISPR/Cas systems of type I, II, and V target the double-stranded DNA of invading phages or conjugative plasmids, but type III and VI target the single-stranded RNA of invading phages, resulting in molecular immunity against invading nucleic acids [8–11].

Using microbial genome data mining and computational prediction approaches to search for previously unexplored class II CRISPR Cas systems has led to the discovery of novel Class II CRISPR systems, including C2c1, C2c2, and C2c3 [7]. C2c1 and C2c3 contain RuvC-like endonucleases similar to Cpf1, and were therefore classified as Class II type V CRISPR/Cas systems, known as type V-B Cas12b and type V-C Cas12c, respectively [12]. In contrast to these other Cas nucleases, Class II candidate 2 (C2c2), designated as Cas13a, has unique features not present in other known Cas proteins. Analysis of the Cas13a protein sequence revealed the presence of two Higher Eukaryotes and Prokaryotes Nucleotide-binding Domains (HEPN), which are exclusively associated with RNase activity [13]. Cas13a contains two Higher Eukaryotes and Prokaryotes Nucleotide-binding domains (HEPN), which are exclusively associated with RNase activity [13]. These distinct characteristics of the Cas13a protein raised the enticing possibility that Cas13a works as a single RNA-guided RNA targeting effector.

A recent work characterizing the functionality of Cas13a of *Leptotrichia shahii* (LshCas13a) has shown that the single effector Cas13a protein is indeed a programmable RNA-guided single-stranded RNA (ssRNA) ribonuclease [11]. Cas13a was shown to provide immunity to a bacteriophage in *Escherichia coli* by interfering with the MS2 lytic ssRNA phage. Cas13a guided by a crRNA containing a 28-nt spacer sequence cleaves target ssRNAs *in vitro* and *in vivo* with protospacer flanking sequence (PFS) of A, U, or C. Cas13a catalytic activity is affected by the secondary structure of the target sequence, where Cas13a tends to preferentially cleave at uracil residues of multiple sites within secondary structures formed in the ssRNA targets. Moreover, the two HEPN domains are responsible for the RNA cleavage activity of Cas13a, and mutation of putative catalytic arginine residues within both domains abolished the RNA cleavage activity, resulting in a catalytically inactive version of the Cas13a enzyme (dCas13a). However, like dCas9, dCas13a retains its ability to bind specifically to the target RNA, and therefore can function as an RNA-guided RNA-binding protein [11, 14]. Cas13a can tolerate single-base mismatches but not double mismatches in the middle of the crRNA spacer sequence–protospacer base paring, pointing to the presence of a central seed sequence. Importantly, the system demonstrated its ability to be reprogrammed to target specific, non-phage RNAs *in vivo* by knocking down the expression of the RFP protein in *E*. *coli*. However, *E. coli* cells expressing RFP and active CRISPR/Cas13a machinery targeting RFP transcript exhibited reduced growth, indicating collateral RNA degradation activity. The collateral Cas13a RNA degradation activities require the successful targeting of the target sequence and the activation of the Cas13a protein. Such collateral targeting could provide a means for engineering programmed cell death for biotechnological applications [11].

East-Seletsky et al. (2016) [15] have shown that Cas13a exhibits dual, distinct functions in processing of the pre-crRNA and degradation of ssRNA targets. These Cas13a activities would facilitate the design and expression of multiple crRNAs under the control of polymerase II promoters for targeting multiple transcripts. Similar to LshCas13a, LbuCas13a from *Leptotrichia buccalis* is activated by a specific crRNA to target a specific ssRNA sequence, but this activation led also to collateral non-specific activities, leading to cell toxicity [15]. The study also showed that the collateral cleavage activity of Cas13a could be advantageous to detect and sense the presence of specific transcripts. Building on this idea, a recent study exploited the collateral effect of the promiscuous RNAse activity of Cas13a upon target recognition to develop a diagnostic tool for *in vitro* detection of RNA, with a single-base mismatch specificity and attomolar sensitivity, demonstrating a wide-range of potential utility in diagnostic and basic research applications [16]. Findings of these studies suggest that Cas13a can function as part of a versatile, RNA-guided RNA-targeting CRISPR/Cas system that holds great potential for precise, robust and scalable RNA-guided RNA-targeting applications.

Plant RNA viruses are responsible for a significant proportion of commercially important plant diseases, infecting a wide range of plant species, and resulting in severe losses in quality and quantity in diverse key crops [17, 18]. Various transgenic strategies based on the pathogen-derived resistance concept have been applied to engineer virus resistance in many plant species [19]. For instance, transgenic plants expressing viral genes (*e.g.* genes encoding viral coat proteins) or RNA sequences of virus origin (to induce RNA interference-mediated resistance) have been shown to confer immunity against viruses from which the genes and RNA sequences were derived [20–22]. In addition, introduction of plant antiviral genes, such as *R* genes, into crop plants has gained prominence as an effective approach to engineer transgenic plants resistant to virus infection [23–26]. Despite the promising potential of these transgenic approaches, many drawbacks have limited their applications in agriculture [27–29].

*Potyviruses* are plant-infecting viruses that belong to the *Potyvirus* genus, one of the largest group of plant viruses besides the *Begomoviruses*. *Potyviruses* represent an economically important group of pathogens capable of infecting and causing a serious damage to a wide range of crop plants. Members of the *Potyviruses*, such as *Turnip mosaic virus* (TuMV), possess long, filamentous particles that are 700–750 nm long, harboring a positive-stranded RNA genome of about 10,000 nucleotides. The single-stranded RNA genome encodes a single long open reading frame (ORF) flanked by terminal untranslated regions. The RNA genome is translated into a single large polyprotein, which is subsequently cleaved by virus-encoded proteinases to yield at least 10 functional proteins [30].

Here, we sought to investigate the possibility of adopting the CRISPR/Cas13a system for interference against RNA viruses in plants and, to that end, we engineered the CRISPR/Cas13a RNA interference system for *in planta* applications. Our study reveals that CRISPR/Cas13a catalytic activities resulted in interference against TuMV-GFP in transient assays and stable overexpression lines of *Nicotiana benthamiana;* and that Cas13a can process long pre-crRNA transcripts into functional crRNAs, resulting in TuMV interference. Thus, our study demonstrates that the CRISPR/Cas13a system is portable to plant species for RNA virus interference, thereby opening myriad possibilities for research and applied uses in plants and other eukaryotic species.

## Results

### Engineering CRISPR/Cas13a machinery for *in planta* expression

Because CRISPR/Cas13a defends bacterial cells against invading RNA viruses, we attempted to test the feasibility and portability of the CRISPR/Cas13a system to provide molecular immunity against RNA viruses in plants. To this end, we selected the potyvirus *Turnip mosaic virus* (TuMV), of the family *Potyvirdiae,* as a target for CRISPR/Cas13a. A recombinant TuMV (TuMV-GFP) expressing GFP was used to provide an easy-to-monitor system for virus infection, replication, and spread, thereby facilitating the assessment of the interference activity of CRISPR/Cas13a system against the RNA virus.

To engineer plants that express the CRISPR/Cas13a machinery, we codon-optimized the *Leptotrichia shahii Cas13a* (*LshCas13a,* WP_018451595.1) nucleotide sequence for expression in plants, and custom synthesized four overlapping fragments to assemble the construct, which we dubbed plant-codon optimized *Cas13a* (*pCas13a*). The four overlapping fragments of *pCas13a* were assembled by unique restriction enzymes to generate a full-length clone flanked by attL1 and attL2 recombination sites, a nuclear localization signal fused to the C-terminus of the protein, and a 3x-HA tag fusion at the N-terminus to facilitate protein detection. Next, *pCas13a* was subcloned into the *pK2GW7* destination binary vector via Gateway recombination reaction to generate *pK2GW7:pCas13a* over-expression clones driven by the *Cauliflower mosaic virus* (CaMV) 35S promoter (Fig. 1A and supplementary Fig.1A). Subsequently, the binary *pCas13a* clone was introduced into *Agrobacterium tumefaciens* via electroporation. Next, Agrobacterium GV3101 cultures harboring the *35S::pCas13a* binary clone were used for transient expression in *N. benthamiana* via agroinfection or for generation of transgenic plants overexpressing the pCas13a protein (pCas13a-OE). We confirmed the pCas13a expression in transient assays in *N. benthamina* leaves and in permanent lines over-expressing pCas13a. Our Western blotting results with anti-HA tag antibody demonstrated that the pCas13a clone was expressed in plants and the correct protein size was detected (Fig. 1B and Supplementary fig. 1B).

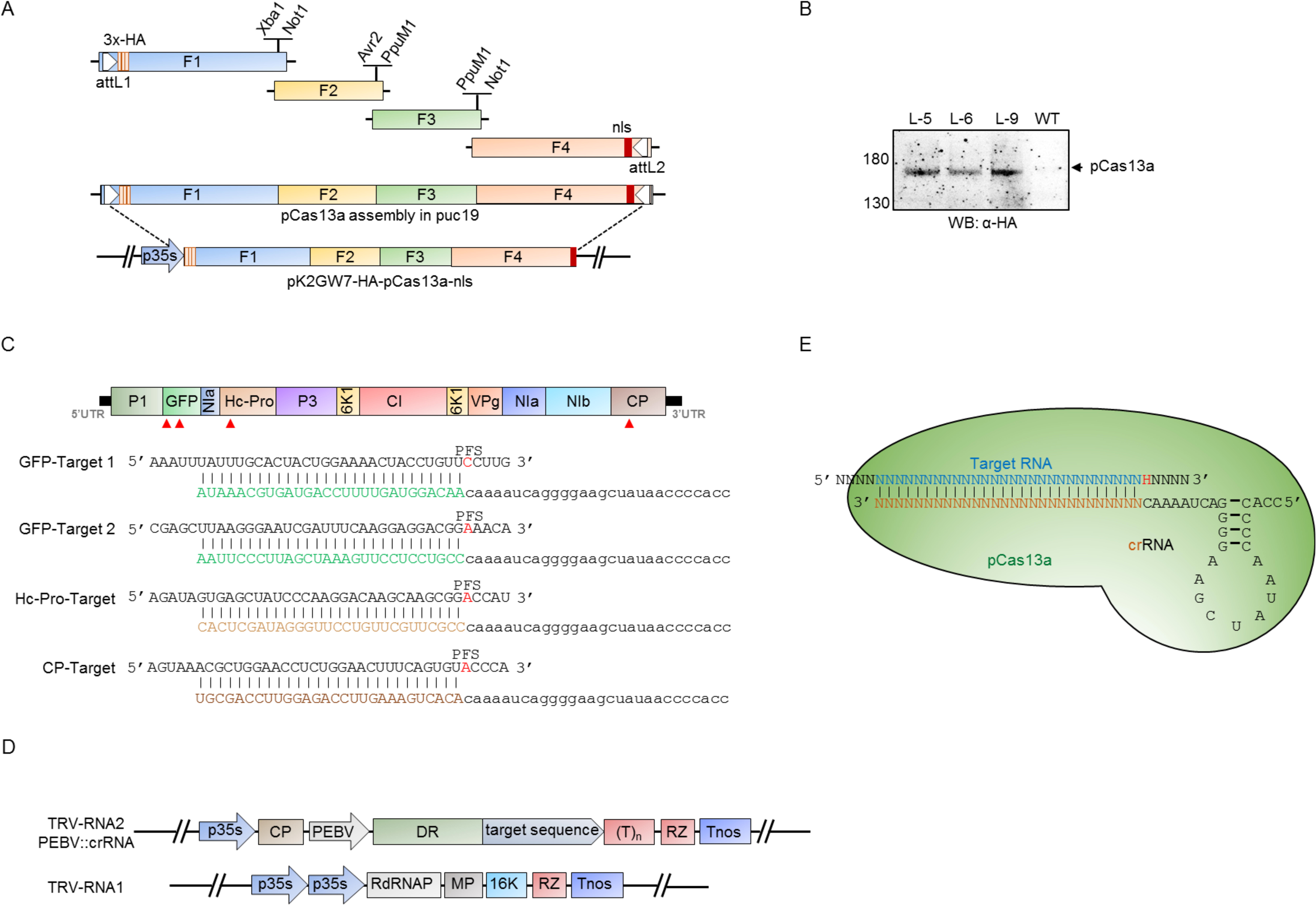

Next, to target the TuMV-GFP virus, we designed and constructed crRNAs complementary to sequences of four different regions of TuMV-GFP genome, including two targets in *GFP* (GFP1 and GFP2), and the helper component proteinase silencing suppressor (*HC-Pro)* and coat protein (*CP*) sequences (Fig.1C). We engineered the RNA2 genome of *tobacco rattle virus* (TRV) to transiently and systemically express crRNAs under the *Pea early browning virus* (PEBV) promoter, as previously described (Fig. 1D) [31]. These crRNAs can be used to test the functionality of CRISPR/pCas13a in transient and stable assays.

### CRISPR/pCas13a interferes with TuMV-GFP *in planta*

To test whether the CRISPR/pCas13a could interfere with TuMV-GFP, we first performed transient assays in *N*. *benthamiana* leaves. Successful interference with the TuMV-GFP genome would result in attenuated replication and spread of the virus, which can be measured by monitoring the level and spread of the virus-mediated GFP expression to systemic leaves during the course of the infection. Mixed *Agrobacterium* cultures carrying the binary *pK2GW7:pCas13a* clones, TRV RNA1, and the engineered TRV RNA2 genome harboring crRNAs against one of the four different TuMV-GFP genome targets (*HC-Pro*, coat protein (*CP*), *GFP* target 1 (*GFP1*), *GFP* target 2 (*GFP2*)), as well as TuMV-GFP infectious clones were co-delivered into *N*. *benthamiana* leaves via agro-infection. In addition, a non-specific crRNA (ns-crRNA) with no sequence similarity to the TuMV-GFP genome was used as a control. The interference activity of the CRISPR/pCas13a system against the TuMV-GFP virus was assessed at 7 days post-infiltration (dpi) by visualizing the GFP signal in the plant systemic leaves under UV light. We observed a reduction of ~50% in the level of GFP signal in the systemic leaves of plants with crRNAs targeting the *HC-Pro* and *GFP2* sequences. In addition, a low, but detectable reduction in the GFP signal was observed with the *CP* and *GFP1* crRNAs compared to the control, whereas no difference in the GFP signal was observed in the wild type *N. benthamiana* plants co-infiltrated with the TRV expressing ns-crRNAs, targeted crRNAs and TuMV-GFP but not *pK2GW7:pCas13a* clones (Supplementary fig. 2A, B). These initial results indicated the functionality and the ability of the CRISPR/pCas13a to interfere with the TuMV-GFP virus *in planta*.

We next assessed the interference activity of CRISPR/Cas13a against the TuMV-GFP in transgenic *N*. *benthamiana* plants constitutively expressing pCas13a protein (pCas13a-OE) under the control of CaMV35S promoter. These transgenic plants were generated via *Agrobacterium*-mediated transformation of *pK2GW7:pCas13a* clones into wild-type *N*. *benthamiana* plants, and the pCas13a protein expression was confirmed by Western blotting (Fig. 1B). The crRNAs targeting the *HC-Pro*, *CP*, *GFP1*, or *GFP2* sequences were delivered into pCas13a-OE plants via the TRV system and expressed under the control of the *PEBV* promoter. In addition, the pCas13-OE plants were co-infiltrated with infectious TuMV-GFP clones via agro-infection. At 7 dpi, the virus-expressed GFP signals were monitored in systemic leaves of the infected pCas13a-OE plants. A substantial reduction, up to 50%, in the GFP intensity was observed with crRNAs targeting the *HC-Pro* and *GFP2* targets. However, consistent with transient assay, crRNAs targeting the *CP* and *GFP1* sequences exhibited only a moderate reduction in the GFP intensity when compared with the ns-crRNA or the empty vector control (Fig. 2A, B).

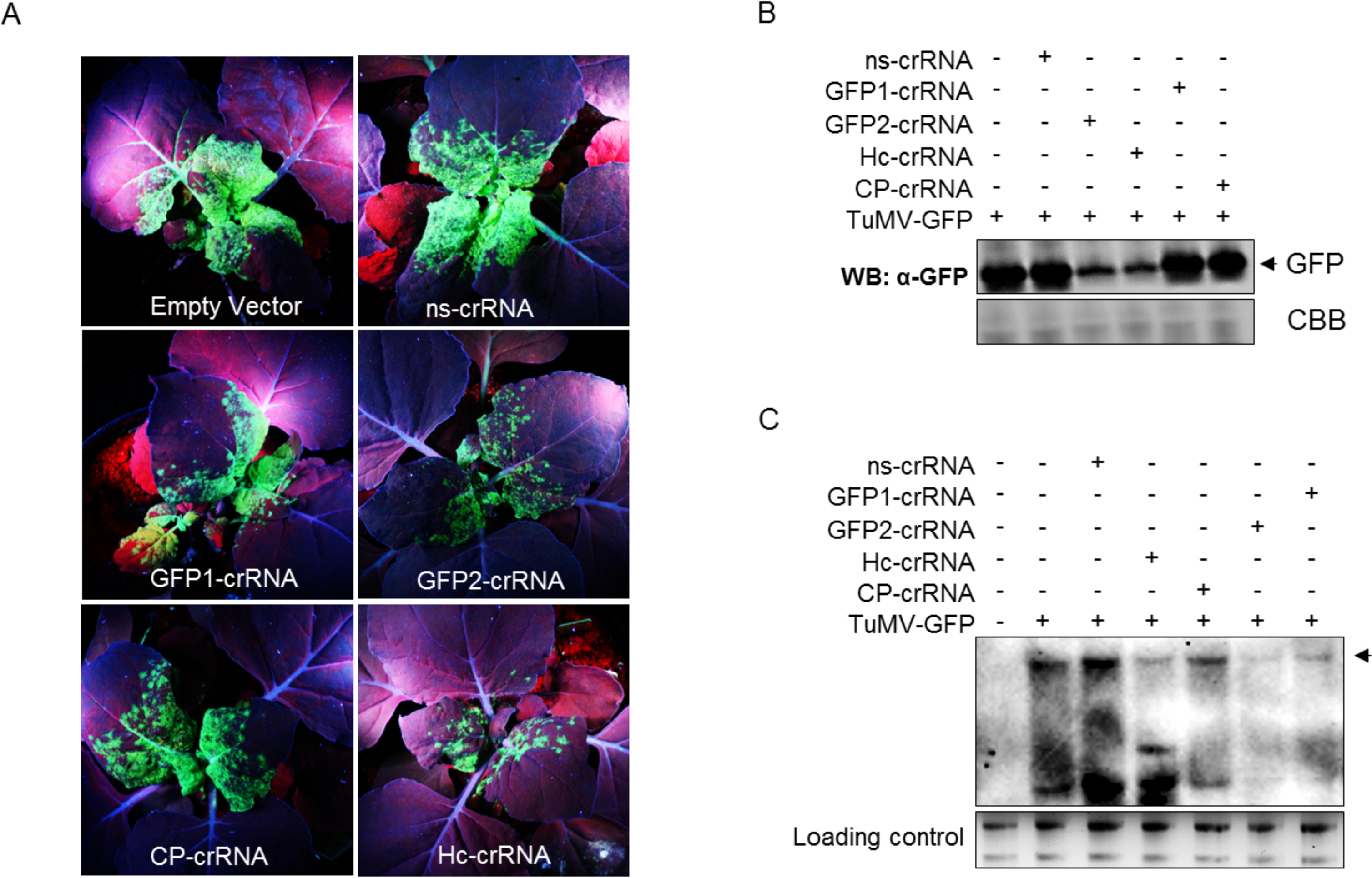

To validate the observed reduction in the GFP signal, Western blotting was used to assess the virus levels in systemic leaves. We found that plants with crRNAs targeting *HC-Pro* or *GFP2* sequences exhibited the largest reduction of GFP levels compared to crRNAs targeting *CP* or *GFP1* target sequences or the controls, corroborating the results obtained from the analysis of the GFP signals (Fig.2C).

To further validate the CRISPR/pCas13a-mediated interference with the RNA genome of TuMV-GFP, total RNA from systemic leaves of the infected plants were isolated, and northern blotting was performed to detect the accumulation of the TuMV genome. Consistent with the level of the GFP protein, the northern blots showed a clear reduction in the accumulation of the TuMV-GFP RNA genome using the crRNAs targeting *HC-Pro* or *GFP2* (Fig. 2D), indicative of targeted degradation of the TuMV-GFP genomic RNA via CRISPR/pCas13a. Importantly, the ns-crRNA showed no interference activity, in contrast to the specific crRNAs, thus validating the specificity and programmability of CRISPR/pCas13a for specific RNA targeting in plants.

### pCas13a processes poly-crRNA into functional crRNAs

Cas13a has been reported to process its own pre-crRNA [15]; therefore, we tested whether pCas13a could process pre-crRNAs in plants. This is of great importance since such ability would enable the generation of multiple functional crRNAs for multiplexed targeting of the same virus, or for simultaneous targeting of multiple RNA and/or DNA viruses in plants. Therefore, we designed CRISPR arrays harboring 28-nt targeting the *HC-Pro*, *GFP1*, and *GFP2* sequences of the TuMV-GFP RNA genome flanked by 28-nt direct repeats (poly-crRNA) (Fig. 3A). In addition, we designed CRISPR arrays harboring non-specific targeting sequences (ns-poly-crRNA). These CRISPR arrays were delivered into wild-type plants (lacking pCas13a) and pCas13a-OE plants via the TRV system to be systematically expressed as long pre-crRNA transcripts, mimicking the endogenous expression of pre-crRNAs in prokaryotes. We analyzed the processing activity of pCas13a by performing northern blotting using probes against the direct repeat regions within the pre-crRNA, with the ultimate goal of detecting the presence of the mature (processed) crRNAs that would indicate the successful generation of mature crRNAs. The northern blotting results showed considerable accumulation of small RNA products in both specific and non-specific poly-crRNAs, consistent with the expected size of the mature crRNAs as compared with the size of the synthetic versions (s-crRNA) of the mature crRNA product (Fig. 3B).

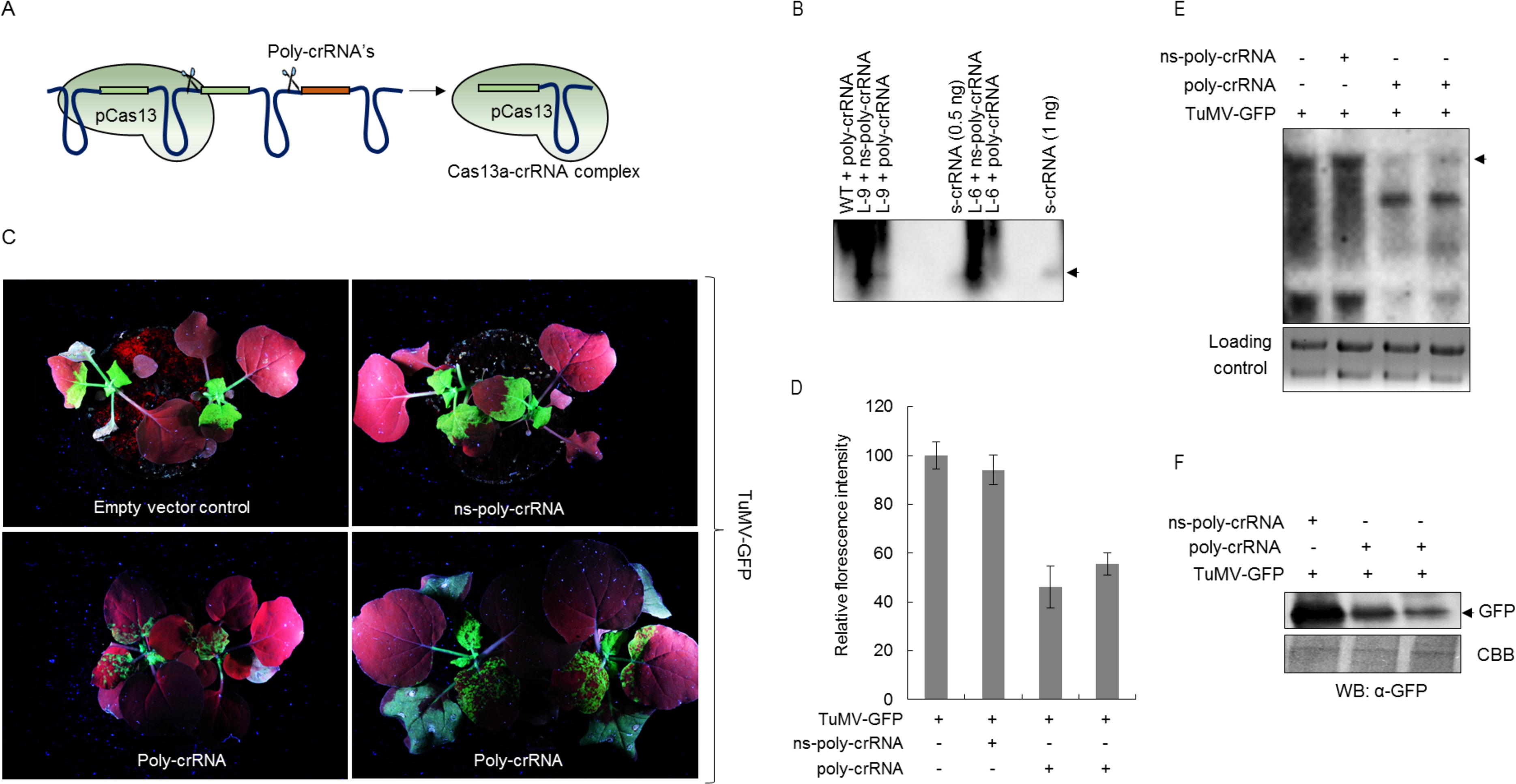

Considering the possibility that the processed “mature” crRNAs would assemble with pCas13a to form active pCas13a–crRNA complexes, we investigated whether the generated mature crRNAs are functional, and therefore, would allow efficient targeting of the TuMV-GFP RNA genome. The pCas13-OE plants were agro-infiltrated with TRV expressing either the poly-crRNA or the ns-poly-crRNAs, and were challenged with the TuMV-GFP virus. The viral GFP signal was determined in systemic leaves at 7 dpi. Strikingly, we detected a significant reduction in the intensity and spread of the viral GFP signal in plants with poly-crRNA. On the other hand, no reduction in the GFP signal was observed with the ns-poly-crRNA relative to the control (Fig. 3C). A quantitative evaluation of the GFP signal revealed an average reduction in the GFP signals of about 50% with the poly-crRNA, whereas no significant reduction was detected with the ns-poly-crRNA (Fig. 3D). Furthermore, results of northern blotting showed a clear reduction in the TuMV-GFP RNA genome level in the case of the poly-crRNA, compared to the undetectable reduction of the TuMV-GFP with the ns-poly-crRNA (Fig. 3E). In addition, western blotting demonstrated a reduction in the protein level of the virus-expressed GFP in plants with the poly-crRNA compared with ns-poly-crRNA, consistent with the results of the GFP signal intensity measurement and northern blot (Fig. 3F). These results indicate the ability of pCas13a to process pre-crRNA transcripts and generate functional crRNAs capable of guiding pCas13a to recognize and interfere with the target RNA virus in plants.

## Discussion

CRISPR/Cas systems provide bacteria and archaea with immunity to fend off invading nucleic acids, and thus provide synthetic bioengineers with rich resources to build systems to edit and regulate the genome and epigenome, and to modulate, modify, and monitor the transcriptome. Therefore, CRISPR/Cas systems can be harnessed for functional biology, biotechnology and genetic medicine. Here, we harnessed CRISPR/Cas13a to selectively interfere with the TuMV RNA virus. Our data demonstrate that CRISPR/Cas13a can mediate molecular interference against a plant RNA virus. Targeting different viral genomic regions for degradation resulted in reduced levels of GFP, coupled with the reduction in the accumulation of the TuMV-GFP RNA genome in systemic leaves. These findings indicate the attenuation of the TuMV virus replication and spread, corroborating the effectiveness of CRISPR/pCas13a in targeting and interfering with the TuMV RNA viruses in plants.

In this study, virus interference was most efficient with crRNAs targeting sequences within the *HC-Pro* and *GFP2* sequences. However, crRNAs targeting the *CP* or *GFP1* target sequences were less effective, suggesting that, among other unexplored factors, secondary structures within the target TuMV-GFP RNA or the presence of RNA binding proteins might influence the accessibility, and thus, the activity of the pCas13a on the target RNA. The better interference activity with the *HC-Pro* crRNA could also be attributed to the fact that TuMV virus uses HC-Pro to suppress the plant’s endogenous antiviral silencing, and any change in HC-Pro level, due to changes in production or turnover, can affect viral replication in the plant cell.

It has been shown that using ≤ 54-nt complementary sequence in TRV system does not activate the post transcriptional gene silencing in plants [32]. However, to exclude the possibility that the 28-nt (spacer sequence of crRNA) can activate the RNAi for the observed TuMV-GFP interference, TuMV-GFP and TRV with crRNAs were co-infiltrated to the wild type *N. benthamiana* plants. Our results (Supplementary fig. 2A lower panel) clearly indicate that the expression of only 28-nt spacers did not lead to any silencing of TuMV-GFP in wild type *N. benthamiana*, confirming that the observed interference is pCas13a dependent.

Catalytically active Cas13a can process long pre-crRNAs *in vitro* and *in vivo*. Importantly *in planta*, the absence of the processed mature crRNA product in the wild-type plants, in contrast to pCas13a-OE plants, provides corroborating evidence that the generation of the mature crRNA is attributed to the presence of pCas13a, consistent with the known role of Cas13a in processing pre-crRNA transcripts [15]. Mature crRNA detection is lower in the presence of its target than the ns-poly-crRNA, probably due to the differential crRNA stability after targeting. Previously different strategies (tRNA and ribozymes) were adopted for multiplex targeting and interference of DNA virus [33, 34]However, the ability of pCas13a to process the poly-crRNA into individual crRNAs permit us to use the pCas13a native potential to produce multi-crRNA from a single poly-crRNA for viral interference. Subsequently, the results obtained from the targeting of TuMV-GFP RNA with a single crRNAs (Fig. 2) and poly-crRNA (Fig. 3 C–F) show analogous patterns of GFP signal, suggesting that the processed crRNA are functional and pCas13a-crRNA complexes are interfering with the TuMV-GFP genome. The observed recovery from virus symptoms in this study shows that the CRISPR/pCas13a system has promise for engineering plants for resistance against plant RNA viruses.

Several questions remain relating to the mechanisms of RNA recognition and subsequent activation of the ribonuclease activities on target and non-target ssRNAs. Liu *et al.* reported that upon target RNA binding and subsequent activation of the ribonuclease activity, Cas13a exhibits indiscriminate degradation of non-target RNAs leading to promiscuous activity[35]. Surprisingly, Cas13a overexpression plants appear similar to the wild-type plants, and introduction of crRNAs with virus target did not lead to cell death. Because promiscuous and collateral RNA-degradation activities may cause cellular toxicity and lead to cell death, our study reveals that such robust promiscuous activity is lacking or too low to be detected in plant cells or eukaryotes in general. While preparing this manuscript, another study about the ability of the Cas13a variants to target mRNA in human cells and plant protoplast was published by Abudayyeh *et al.* (2017) [36]observing no promiscuous activity of Cas13a in eukaryotic cells. Further studies are needed using different Cas13a variants and exhaustive crRNAs against viral and endogenous RNA targets. Future work would focus on Cas13a variants that exhibit robust ribonuclease activity. Alternatively, the RNA targeting and binding functions could be used for selective degradation of targets through the use of other robust ribonuclease domains fused to the Cas13a backbone. Our findings show that Cas13a is an efficient RNA-guided ribonuclease that can be programmed to target and degrade viral RNA genomes, thereby providing a promising and an effective tool for a variety of RNA manipulations and specifically interference against RNA viruses in plants, and in eukaryotic cells in general.

## Material and Methods

### Design and construction of CRISPR/Cas13 machinery for in planta expression

We designed a pCas13a protein including 3x-HA, NLS, attL1, and attL2 sites. The nucleotide sequence of *pCas13* was designed as four overlapping fragments with unique restriction enzymes to facilitate sequential cloning and assembly into a single fragment. These four fragments were custom synthesized into the pUC19(-MCS) plasmid by Blue Heron Biotech gene synthesis services (Blue Heron Biotech, Bothell, WA, USA). The first fragment (fragment 1) is preceded by the attL1 sequence and the last fragment (F4) ends with the attL2 sequence for subsequent LR Gateway recombination reaction. To assemble the four fragments into a single fragment, fragments 1 and 2 were assembled by restriction digestion with Xba1 and PpuM1 enzymes and fragment 3 and 4 were assembled with pPuM1 and Not1 to generate pUC19- (F1+F2) and pUC19-(F3+F4), respectively. Subsequently, the pUC19-F3+F4 clone was sub-cloned into pUC19-F1+F2 using Avr2 and NotI enzymes to generate a single clone with the complete and in-frame sequence of *pCas13a*. The *pUC19-pCas13a* construct was confirmed by Sanger sequencing using overlapping primers. Next, we sub-cloned *pCas13a* into the *pK2GW7* binary vector using LR Gateway recombination cloning to generate pK2GW7-pCas13a for expression in plants under the control of the 35S promoter.

The crRNAs were designed as primer dimers with overhang and were cloned under PEBV promoter in TRV RNA2 by XbaI and BamHI.

The poly-crRNA and ns-poly-crRNA were custom synthesized in pUC19 with unique sites (BamHI and SacI). Both the poly-crRNA and the ns-poly-crRNAs were cloned to the TRV, RNA2 by BamHI and SacI.

## Plant Material

*N. benthamiana* plants expressing Cas13a with and without crRNA were regenerated using the previously developed protocol [31]. The *pK2GW7-pCas13a* construct was used to generate plants (*N. benthamiana* or resistant) for programming resistance to TuMV. All plants were selected on 50 mg kanamycin in ½ Murashige-Skoog agar media.

## Agro-infiltration of *N. benthamiana* leaves and GFP imaging

Constructs, harboring the TuMV-GFP infectious clone, TRV RNA2 empty or containing crRNA under PEBV promoter, TRV RNA1, were individually electroporated into *Agrobacterium tumefaciens* strain GV3101. Overnight-grown single colonies in selective medium were centrifuge and suspended, in infiltration medium (10 mM MES [pH 5.7], 10 mM CaCl_2_, and 200 µM acetosyringone), and incubated at ambient temperature for 2 h. For infiltration into Cas13a-OE plants, cultures were mixed at an OD_600_ ratio of 0.05:1:1 (TuMV-GFP, TRV RNA2, TRV RNA1) and infiltrated into 3- to 4-week-old leaves of *N. benthamiana* Cas13a-OE plants with a 1-mL needleless syringe. GFP expression was observed at 3, 7, and 10 dpi using a hand-held UV light. Photos were taken with a Nikon camera both in normal light and under UV light. GFP signal quantifications were done using ImageJ software. Leaves samples were collected at 7 and 15 dpi for molecular analyses.

### Immunoblot Analysis

Total proteins were extracted from 100 mg of sample using extraction buffer (100 mM Tris-Cl pH8, 150 mM NaCl, 0.6% IGEPAL, 1 mM EDTA, 3 mM DTT with protease inhibitors, PMSF, leupeptin, aprotinin, pepstatin, antipain, chymostatin, Na_2_VO_3_, NaF, MG132, and MG115. Proteins were separated on a 10% polyacrylamide gel. Immunoblot analysis was carried out using mouse α-GFP (1:2000; Invitrogen) for TuMV GFP and rat α-HA (1:500) antibody for pCas13a. The antigens were detected by chemiluminescence using an ECL-detecting reagent (Thermo Scientific)

### RNA isolation and northern blot analysis

Total RNA was extracted from virus-infected plants using the Direct-zol RNA MiniPrep Plus (Zymo Research) according to the manufacturer’s recommendations. For each sample, 10–15 μg of RNA was separated on a denaturing 2% agarose gel, blotted on a Hybond-N^+^ (GE Healthcare) membrane and hybridized with a DIG-labelled probe. For virus expression analysis, a DIG-labelled RNA probe was synthesized using DIG Northern Starter Kit (Roche) and manufacturer’s instructions were followed. For crRNA detection, 5´-end DIG-labelled oligonucleotide (IDT) was used. DIG application manual (Roche) was followed for capillary transfer, hybridization and detection. Northern blots were repeated in three independent experiments with the same results.

## Acknowledgements

We wish to thank Professor James Carrington for TuMV-GFP and members of the Laboratory for Genome Engineering at King Abdullah University of Science and Technology for helpful discussions and comments on the manuscript. This publication is based upon work supported by the King Abdullah University of Science and Technology (KAUST) Office of Sponsored Research (OSR) under Award No. OSR-2015-CRG4-2647.

## Author Contributions

MM conceived research; MM, RA, ZA, HB, AM, and SWD designed research; RA, ZA, HB, FA, MZK, and AM performed research; MM, RA, ZA, HB, AM, and SWD analyzed data; MM, AM, ZA, and MM wrote the paper.

